# A Chemical Mutagenesis Approach to Insert Post-Translational Modifications in Aggregation-Prone Proteins

**DOI:** 10.1101/2022.01.28.478135

**Authors:** Ying Ge, Athina Masoura, Jingzhou Yang, Francesco A. Aprile

## Abstract

Neurodegenerative diseases are a class of disorders linked to the formation in the nervous system of fibrillar protein aggregates called amyloids. This aggregation process is affected by a variety of post-translational modifications, whose specific mechanisms are not fully understood yet. Emerging chemical mutagenesis technology is currently striving to address the challenge of introducing protein post-translational modifications, while maintaining proteins stable and soluble during the modification reaction. Several amyloidogenic proteins are highly aggregation-prone, and current modification procedures lead to unexpected precipitation of these proteins, affecting their yield and downstream characterization. Here, we present a method for maintaining amyloidogenic proteins soluble during chemical mutagenesis. As proof-of-principle, we applied our method to mimic the phosphorylation of the serine 26 and the acetylation of the lysine 28 of the 40-residue long variant of amyloid-β peptide, whose aggregation is linked to Alzheimer’s disease.

## Introduction

Dementia is an umbrella term that refers to several pathologies characterized by progressive and irreversible damages to the nervous system. They are a major cause of morbidity and mortality across the globe, with over 78 million people estimated to be affected by 2030 worldwide.^1^ Alzheimer’s disease (AD), the most common form of dementia, is distinguished by disease hallmarks including amyloid-β (Aβ) plaques and tau-containing neurofibrillary tangles.^2,3^ Aβ is an aggregation-prone peptide produced from aberrant cleavage of the amyloid precursor protein (APP), an integral membrane protein abundant at the synapses, by the sequential proteolytic cleavage of the β- and γ-secretases.^4^ Genetic mutations of APP and of the secretase genes promoting this proteolytic pathway and are linked to familial AD.^5–7^ Furthermore, most cases of AD are sporadic and post-translational modifications (PTMs) can be induced by environmental factors such as inflammation.^8^ PTMs including truncation, serine phosphorylation, lysine acetylation, methionine oxidation and polyglutamylation are found in Aβ aggregates, with reported effects such as increasing or reducing the rate of aggregation and plaque formation.^2,8^ However, detailed biomolecular and mechanistic studies of each PTM is hindered by the lack of site-specific tools to introduce them. To date, PTMs in Aβ have been generated by solid-state peptide synthesis,^9–11^ enzymatic reaction,^12,13^ and non-site specific chemical modification.^14^ However, these methods have their drawbacks. Solid-state synthesis is costly. Enzymatic reactions are limited in scope, whilst chemical modifications tend to react with multiple residues of the same type. In contrast, site-specificity and product versatility can be combined using a dehydroalanine (Dha)-based method of introducing PTM mimetics developed by Davis and co-workers.^15,16^ Briefly, cysteine is converted to Dha via an alkylation-elimination reaction with a dibromide compound, allowing subsequent Michael addition with a variety of thiol-containing compounds. As cysteine can be introduced via molecular cloning techniques, installation of PTMs can be carried out in a site-specific yet versatile manner.

This method has been applied to histone H3,^15,16^ as well as to a single-domain antibody and the amyloidogenic protein Tau.^17,18^. However, additional challenges are faced in the modification of Aβ due to its hydrophobic and aggregation-prone nature. Procedures required for the chemical reactions, such as heating and shaking, could lead to protein precipitation and aggregation, thus reducing the yield and limiting downstream application. Here, we establish a facile, high-yield protocol of introducing PTM mimetics in Aβ, while maintaining the peptide soluble, by utilizing a solubility-enhancing domain.^19^ As a proof of concept, we installed mimetics of two PTMs: phosphorylation of S26 and acetylation of K28. K28 has been reported to form salt bridge with D23 in Aβ40,^11,20^ and with the carboxyl group of A42 in Aβ42,^21^ although K28 acetylation only had a limited effect on the aggregation of Aβ42.^10^ In contrast, S26 phosphorylation in Aβ40, which results in an alternative pS26-K28 salt bridge, appears to promote the accumulation of cytotoxic oligomeric species while reducing the formation of fibrils, leading to a reduction of fluorescence intensity in Thioflavin T (ThT) and Congo red assays^11,13^ By successfully introducing mimetics of these two PTMs, we overcome the unique difficulties of carrying out chemical biology on a peptide with a high propensity to aggregate. We also demonstrate that the aggregation behaviors of these peptides agree with previous reports.^10,11,13^ This method can be readily adapted to introduce other types of PTMs in Aβ and enable a wide range of biophysical studies.

## Results and Discussion

### Experimental Strategy

Aβ40 and its cysteine variants are expressed in *Escherichia coli* (*E. coli*) as a fusion protein with spider silk protein domain (SD) as previously reported.^19^ Purification and modification are carried out on the fusion protein (Figure 1), as we theorize that enhanced solubility afforded by the SD would allow multiple steps of chemical modification to be carried out on Aβ. First, SD-Aβ40 (or variant) is purified under reducing condition using affinity chromatography, via its N-terminal His6 tag. As there is no cysteine present in Aβ40 natively, the introduced cysteine can be site-specifically converted to Dha by reacting with of 2,5-Dibromohexanediamide (DBHDA). Dha26 is subsequently converted to phosphoserine mimetic phosphocysteine (pC) by reaction with sodium thiophosphate, while Dha28 is reacted with N-acetylcysteamine to mimic acetyllysine. The resulting residue is hereafter referred to as Ac(S)K28 as the sidechain Cβ of lysine is substituted with S. Finally, the addition of Tobacco Etch Virus (TEV) protease separates the SD from Aβ, and size-exclusion chromatography (SEC) enables the removal of SD to obtain modified Aβ.

**Figure 1.**
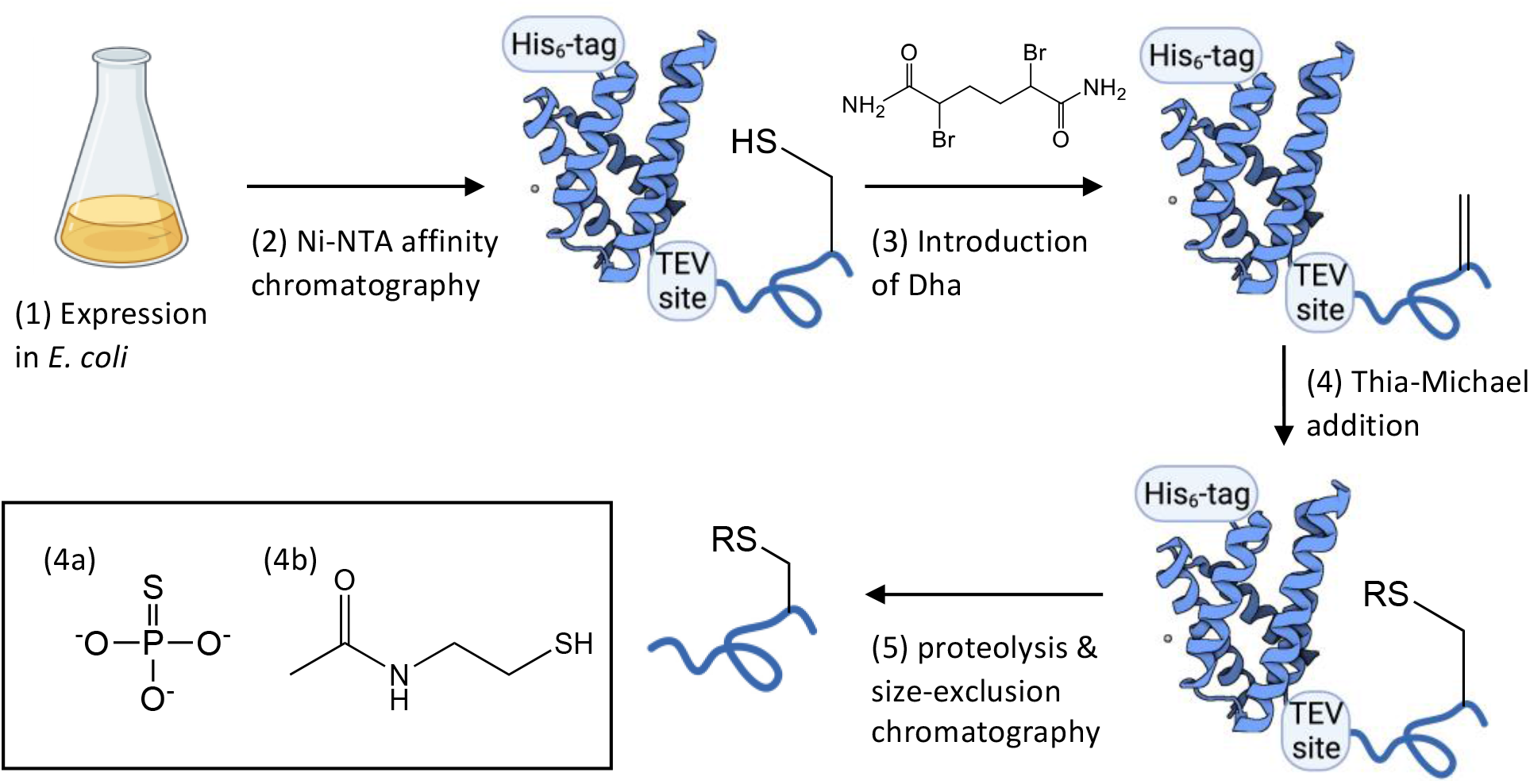
Experimental Strategy. (1) SD-Aβ40 with a single cysteine is expressed as a His_6_-tagged fusion protein in *E. coli* BL21(DE3) cells. (2) The fusion protein is purified from the cell lysate by Ni-NTA affinity chromatography. (3) Purified fusion protein is reacted with DBHDA to convert the cysteine residue into a Dha. (4) Dha is then converted into phosphocysteine or acetyllysine mimic by thia-Michael addition with sodium thiophosphate (4a, R = PO_3_) or N-acetylcysteamine (4b, R= C_4_H_8_NO), respectively. (5) The SD is removed via proteolysis by the addition of TEV protease, and the digestion mixture was subjected to size-exclusion chromatography to remove the SD and TEV, and to obtain monomeric Aβ.

### Introduction of Dha and PTM mimetics to SD-Aβ40

SD-Aβ40-S26C and SD-Aβ40-K28C were purified as a fusion protein by Ni-NTA affinity chromatography. Mass spectrometry (MS) confirmed the serine/lysine to cysteine mutations, as the observed mass matches that of the expected fusion protein minus an N-terminal methionine, which is thought to have been lost during purification or MS analysis (Figure 2A, 2E and S1, S5). In SD-Aβ40-K28C, an additional peak with a +75 Da mass shift was observed (Figure 2E and S5), which may be a BME-adduct resulted from the purification process, and disappears after modification with DBHDA (Figure 2F and S6). A mass shift of −34 Da was observed (Figure 2B, 2F and S2, S6) following incubation of the protein with DBHDA at 37 °C for 3 h, indicating the conversion of the cysteine into Dha. For the final modification step, SD-Aβ40-Dha26 was reacted with sodium thiophosphate, resulting in a mass shift of +80 Da compared to SD-Aβ40-S26C, (Figure 2C and S3) in accordance with the formation of pC. For SD-Aβ40-Dha28, the formation of an acetyllysine mimic was observed after incubation with N-acetylcysteamine (Figure 2G and S7).

**Figure 2.**
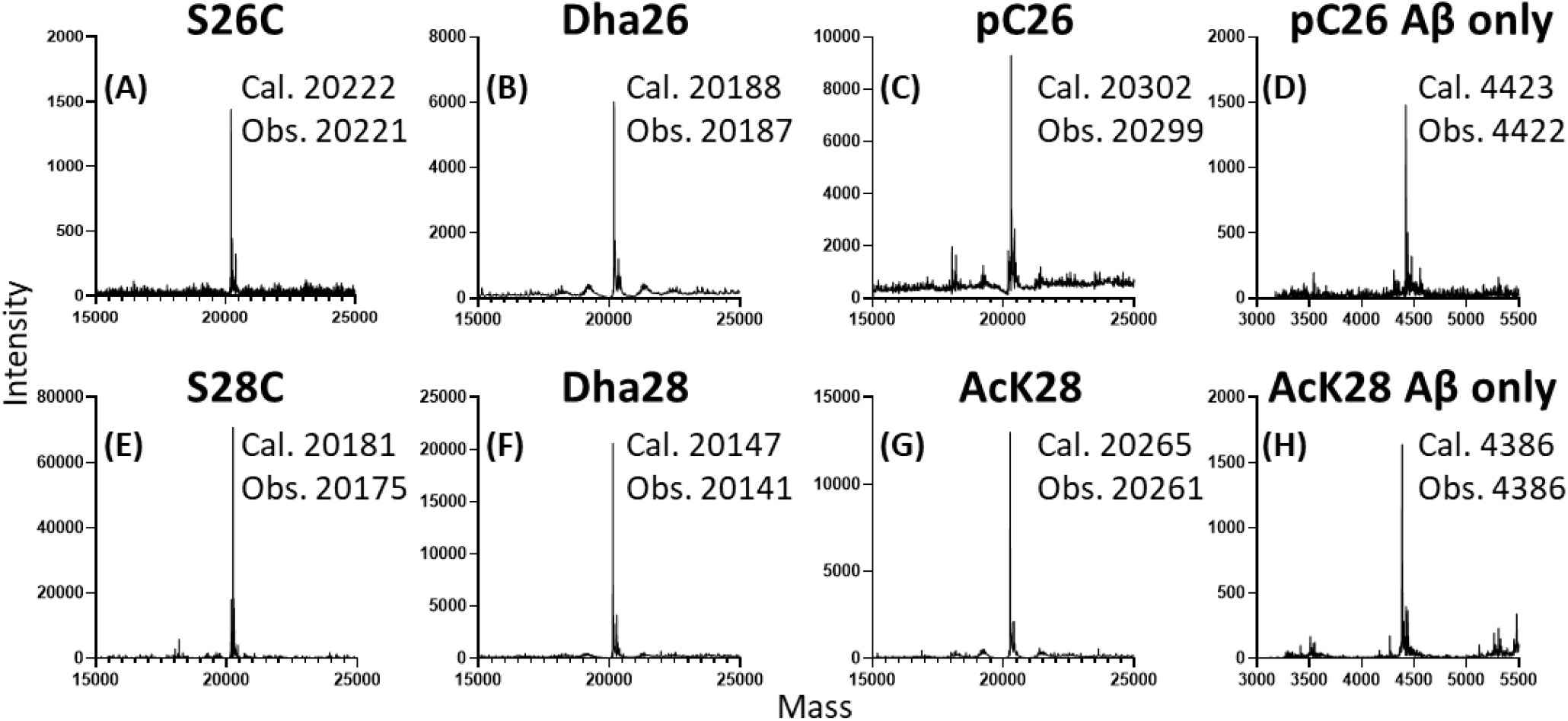
Chemical mutagenesis of SD-Aβ40. (A-C) Deconvoluted mass spectra of SD-Aβ40-S26C with C26 (A), Dha26 (B) and pC26 (C), showing calculated and observed mass of each variant. (D) Deconvoluted mass spectrum of Aβ40-pC26 after TEV cleavage and SEC purification. (E-G) Deconvoluted mass spectra of SD-Aβ40-K28C with C28 (E), Dha28 (F) and Ac(S)K28 (G). (H) Deconvoluted mass spectrum of Aβ40-AcK28 after TEV cleavage and SEC purification. Cal., calculated; Obs., observed.

### Purification of Monomeric Aβ40 variants

To produce authentic (i.e., with no overhanging initial methionine) Aβ40 peptide without the SD, the fusion protein was incubated with TEV protease to remove the N-terminal silk domain (Figure 1, Figure 3A). The resulting peptide begins with an aspartic acid, same as naturally occurring Aβ peptides. The proteolyzed sample was subjected to SEC and fractions containing purified Aβ peptides are collected for subsequent characterization (Figure 3). MS analysis shows Aβ40 peptide at the expected mass with high homogeneity (Figure 2D, 2H and S4, S8), confirming that the modifications remain stable after removal of the silk domain by TEV protease cleavage and the SEC process.

**Figure 3.**
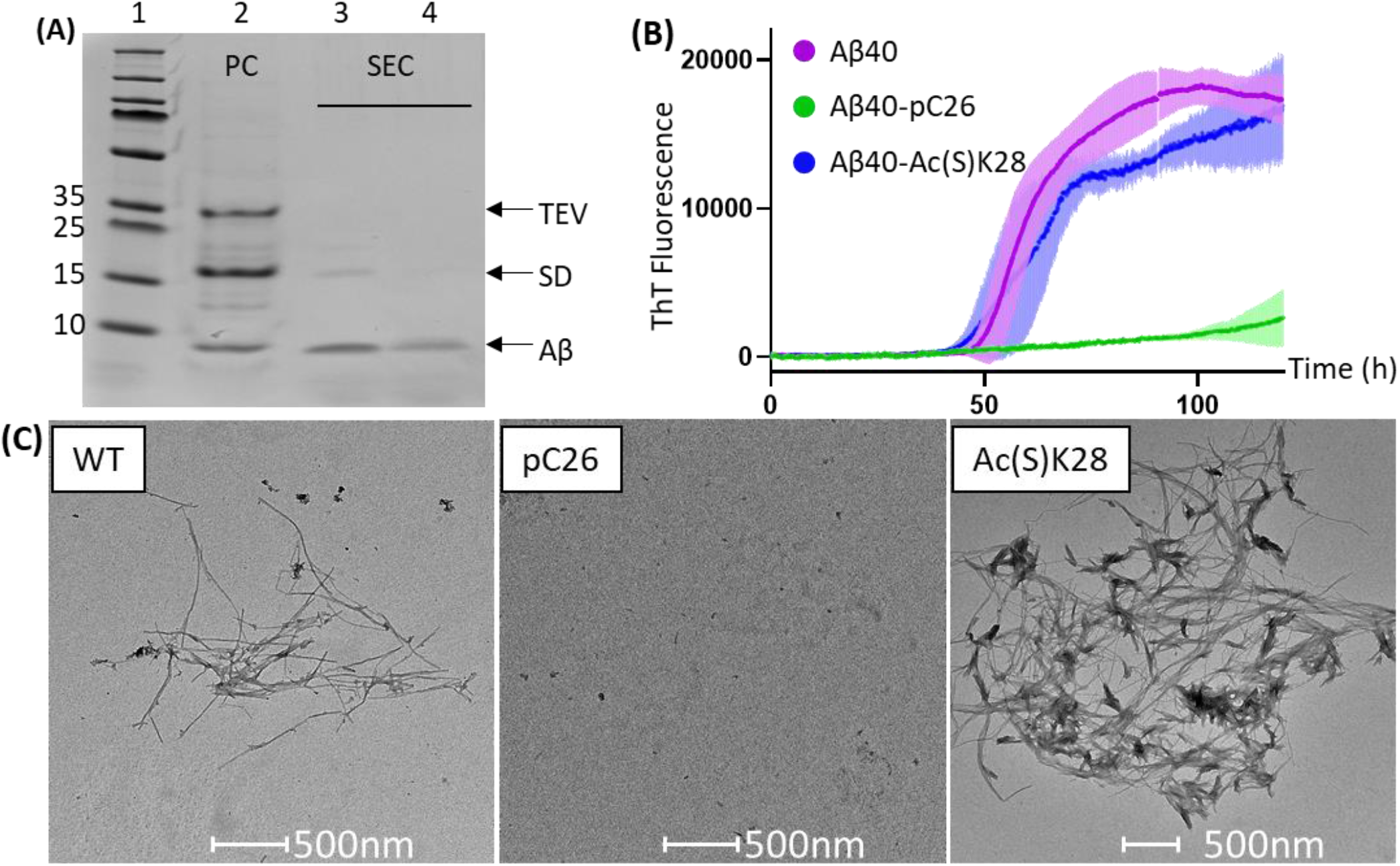
(A) Representative SDS-PAGE showing the purification of Aβ40 (or PTM variants) from the fusion protein-TEV reaction mix. Lane 2, SD-Aβ40 TEV reaction mixture (PC, pre-column). Following each SEC purification multiple fractions are collected and the purest (in this case lane 4) are used in aggregation assays. (B) ThT aggregation assay of 5 μM recombinant Aβ40 (magenta), Aβ40-pC26 (green) or Aβ40-Ac(S)K28 (blue) peptides, and. Fluorescence of 20 μM ThT in buffer alone was recorded in parallel and subtracted as baseline. Error bars show standard deviation (N=5). Inset shows the aggregation of Aβ40-pC26. (C) TEM images showing long, thin fibrils observed in Aβ40, no fibrils in Aβ40-pC26, and shorter, clustered fibrils in Aβ40-Ac(S)K28 after 163 h of incubation at 37 °C.

### Comparison of the aggregation behaviors of Aβ40, Aβ40-pC26 and Aβ40-Ac(S)K28

ThT assays were carried out to compare the aggregations of Aβ40, Aβ40-pC26 and Aβ40-Ac(S)K28 to determine whether our modifications have consistent effects with the PTMs of which they are mimetics. The ThT kinetic of Aβ40-Ac(S)K28 was slower but overall comparable to that of Aβ40 (Figure 3B). This observation agrees with a previous study of Aβ42, where K28 acetylation had limited effect on the protein aggregation rate.^10^ Transmission electron microscopy (TEM) on samples taken at the plateau of the aggregation showed long, thin fibrils for Aβ40 (Figure 3C). Shorter, heavily clustered fibrils were observed for Aβ40-Ac(S)K28 (Figure 3B), suggesting that K28 acetylation may affect the morphology of aggregates. In contrast, Aβ40-pC26 showed a slow, close-to-linear increase in ThT fluorescence (Figure 3B), similar to solid-phase synthesized Aβ40-pS26.^11,13^ A lack of fibrillar aggregates observable by TEM in Aβ40-pC26 samples after incubation (Figure 3C) confirms the disruption to fibrillization caused by phosphorylation of residue 26.

The current study is a proof of concept of a strategy to post-translationally modify in a site-specific manner aggregation-prone proteins. Although the final products are likely an epimeric mixture due to the planar conformation of Dha, several works have demonstrated the suitability of these PTM mimetics as research tools. Histone H3 containing pC generated via the Dha intermediate were shown to respond to pS-specific antibodies, and to be substrates of phosphatases.^16^ Installing pC at S356 of tau results in the inhibition of tubulin polymerization, an effect also observed when S356 is phosphorylated by other methods.^18^ Our results further demonstrated that PTMs mimetics introduced via the Dha-intermediate can be used to study the effect of the corresponding PTMs on protein aggregation.

## Methods

### Cloning, Protein Expression and Purification of the Silk Domain Aβ40 Fusion Proteins

The mutations S26C and K28C were introduced to SD-Aβ40 fusion protein^19^ via site-directed mutagenesis using a QuikChange Lightning Kit (Agilent, USA) following a modified protocol.^22^ The successful introduction of mutations was confirmed by sequencing (Genewiz). Chemically competent *E. coli* BL21(DE3) cells were transformed with the wildtype or mutated plasmids and used to overexpress proteins according to established protocol.^19^ Briefly, cells were cultured in LB-kanamycin medium at 37°C, 120 rpm until OD600nm reached 0.8-0.9, when the temperature was lowered to 20 °C and expression was induced by the addition of 0.1mM Isopropyl β-d-1-thiogalactopyranoside (IPTG). After incubation overnight, cells were harvested by centrifugation at 6000 g for 15 min, and the pellets were frozen and thawed prior to re-suspension in binding buffer (8 M urea, 20 mM Tris-HCl pH 8.0, 15 mM imidazole) and sonication at 500 W, 20% amplitude for 10 min (15 s on, 45 s off). Cell lysate was cleared by centrifugation at 38,758 g and the supernatant was passed through a syringe filter with a diameter of 0.45 μm. The filtrate was applied to a HisTrap HP column (Cytiva) and the target protein was eluted with a mix of 40% binding buffer and 60% elution buffer (8 M urea, 20 mM Tris-HCl pH 8.0, 300 mM imidazole). The elute was collected and dialyzed against 50 mM phosphate buffer pH 8.0 for 16 h. For cysteine-containing variants, binding and elution buffers were supplemented with 3 mM β-mercaptoethanol (BME). Protein concentrations were calculated using A280 nm values measured on a Nanodrop (Thermo Scientific) and extinction co-efficient values obtained from the Expasy ProtParam tool^23^.

### Introduction of Dha

2 mM tris(2-carboxyethyl)phosphine (TCEP) and 400 molar equivalent of DBHDA was added to 200 μM SD-Aβ40-S26C or SD-Aβ40-K28C, and the reaction was incubated at 37 °C with shaking at 400 rpm for 3 h. After the precipitate was removed by centrifugation, small molecules were removed by passing the sample through a HiTrap desalting column with Sephadex G-25 resin (Cytiva) pre-equilibrated in 0.1 M sodium phosphate buffer pH 8.0, or by dialysis. For mass spectrometry (MS) analysis (liquid chromatography electrospray, LC-ES), the samples were desalted using Zeba Spin columns (Thermo Scientific) with 7K MWCO.

### Generation of phosphocysteine

480 mg sodium thiophosphate (NaSPO_3_) was dissolved in 186 μl H_2_O and 200 μl 5 M HCl. This stock was added to 100 μM Dha-containing protein (SD-Aβ40-Dha26) at 1:5 volume ratio. Reactions were carried out at 37 °C with shaking at 400 rpm for 6 h in the presence of 1.5 M urea. Excess salt was removed by dialysis in 0.1 M sodium phosphate buffer pH 8.0 prior to TEV cleavage. For MS analysis, samples were desalted using Zeba Spin columns (Thermo Scientific) with 7K MWCO.

### Generation of acetyllysine mimic

50 μl of N-Acetylcysteamine were added to 1.2 ml of 100 μM SD-Aβ40-Dha28. Reactions were incubated at room temperature (25 °C) with shaking at 400 rpm for 3 h. Excess salt was removed by dialysis in 0.1 M sodium phosphate buffer pH 8.0 prior to TEV cleavage. For MS analysis, samples were desalted using Zeba Spin columns (Thermo Scientific) with 7K MWCO.

### Removal of the SD and Purification of the Aβ40 Variants

SD-Aβ40 and variants were dialyzed against 0.1 M phosphate buffer pH 8.0 and the SD was cleaved from Aβ40 by incubating with TEV protease at 20:1 molar ratio for 1 h at room temperature followed by 23 h at 4 °C. After proteolysis the sample was dissolved in 6 M guanidine-HCl and subjected to size exclusion chromatography on a Superdex 75 increase 10/300 GL column (Cytiva) pre-equilibrated in 20 mM phosphate buffer pH 8.0 supplemented with 200 μM EDTA, to separate the Aβ40 peptide from TEV and SD. Fractions were analyzed by SDS-PAGE and only those containing pure Aβ were used for subsequent applications. Concentration is calculated from the UV absorption reading on an ÄKTA Pure protein purification system operated with the Unicorn 7 software. For MS analysis, the samples were buffer exchanged into H_2_O using an Amicon concentrator with 3K MWCO (Millipore).

### ThT Aggregation Assay

Aβ40 and PTM variants were mixed and incubated with 20 μM ThT and 0.02% sodium azide at 37 °C without shaking in 20 mM sodium phosphate buffer, pH 8.0, 200 μM EDTA, in a black 96-well non-binding microplate with clear bottom (Greiner #655906). Fluorescence was monitored in a CLARIOstar Plus plate reader with excitation filter of 440±10 nm and emission filter of 480±10 nm.

### Transmission Electron Microscopy

Aβ40 and PTM variants were mixed and incubated with 20 μM ThT and 0.02% sodium azide at 37 °C without shaking in 20 mM sodium phosphate buffer, pH 8.0, 200 μM EDTA. Samples were collected at 163 h of the incubation and deposited onto carbon-coated copper mesh grids and negatively stained with 2% (w/v) uranyl acetate. The samples were then viewed with a FEI Tecnai T12 Spirit 120 kV transmission electron microscope.

## AUTHOR INFORMATION

### Author Contributions

Y.G and F.A.A designed the research. Y.G, A.M and J.Y performed the experiments. Y.G, A.M, J.Y and F.A.A analyzed the data. Y.G., A.M and F.A.A. wrote the manuscript.

### Funding Sources

We thank UK Research and Innovation (Future Leaders Fellowship MR/S033947/1), the Alzheimer’s Society, UK (Grant 511), and Alzheimer’s Research UK (ARUK-PG2019B-020) for support.

### Notes

The authors declare no competing financial interest.

## ACKNOWLEDGMENT

MS analyses were carried out by Dr Lisa Haigh of the Chemistry Mass Spectrometry Facility, Imperial College London. We thank Dr Henrik Biverstål (Karolinska Institutet) for providing the expression plasmid for the silk domain-Aβ40 fusion protein.

## ABBREVIATIONS

Aβ: amyloid-β
AcK: acetyllysine
Ac(S)K: thiol-containing mimic of AcK
AD: Alzheimer’s disease
APP: amyloid precursor protein
BME: β-mercaptoethanol
DBHDA: 2,5-dibromohexanediamide
DTT: dithiothreitol
EDTA: ethylenediaminetetraacetic acid
IPTG: isopropyl β-D-1-thiogalactopyranoside
LC-ES: liquid chromatography electrospray
MS: mass spectrometry
pC: phosphocysteine
pS: phosphoserine
PTM: post-translational modification
SD: silk domain
SDS-PAGE: sodium dodecyl sulphate–polyacrylamide gel electrophoresis
SEC: size-exclusion chromatography
TCEP: tris(2-carboxyethyl)phosphine
TEM: Transmission electron microscopy
TEV: Tobacco Etch Virus
ThT: Thioflavin T

## Notes

### Competing Interest Statement

The authors have declared no competing interest.

