## Supplemental information for "A Chemical Mutagenesis Approach to Insert Post-Translational Modifications in Aggregation-Prone Proteins"

**Supplementary information includes:**

- Scheme S1
- Figures S1-S8

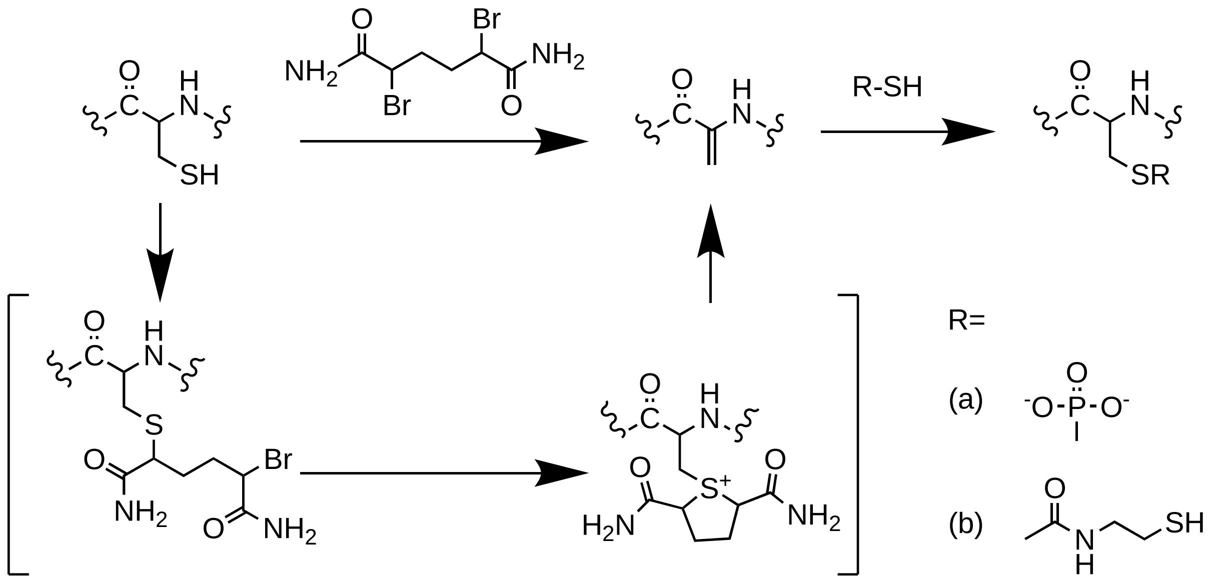


Scheme S1 Reaction mechiansm of PTM installation via Dha.

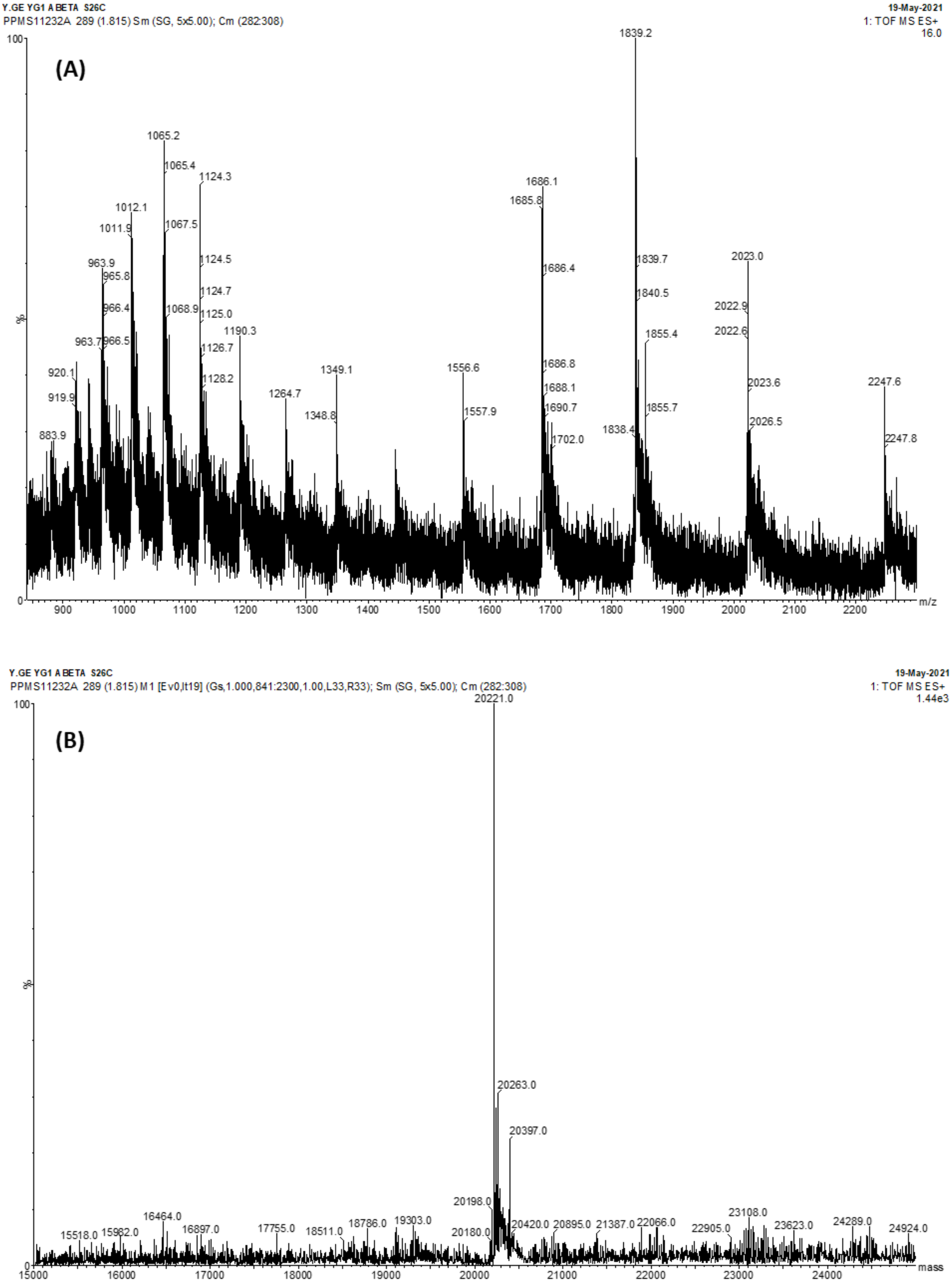


Figure S1. LC-ES mass spectra (A) and deconvolution data (B) of Aβ40 S26C as fusion protein. The calculated mass (minus the N-terminal methionine) is 20222 and the observed deconvoluted mass is 20221.

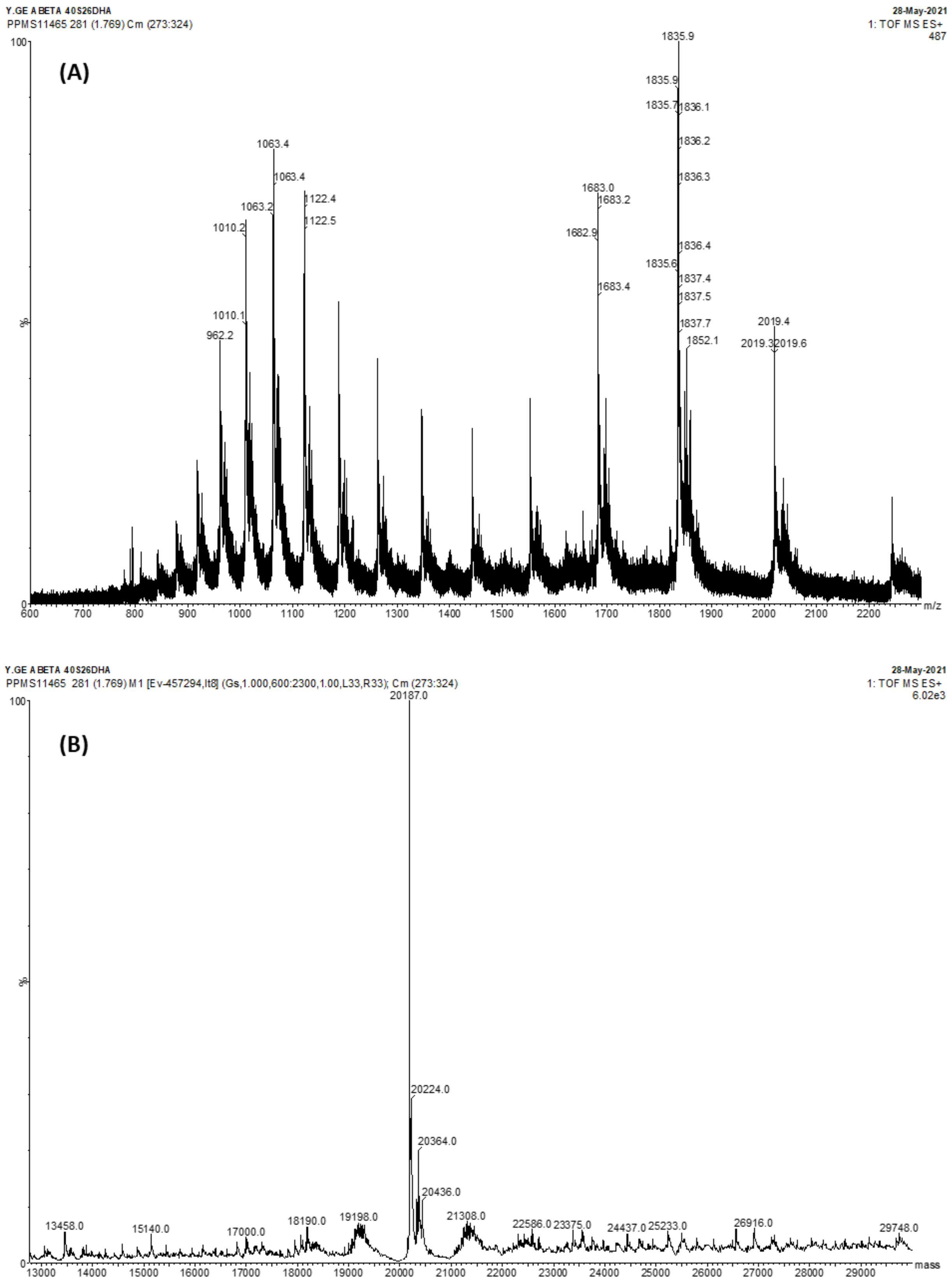


Figure S2. LC-ES mass spectra (A) and deconvolution data (B) of SD-Aβ40-S26Dha. The calculated mass (minus the N-terminal methionine) is 20188 and the observed deconvoluted mass is 20187.

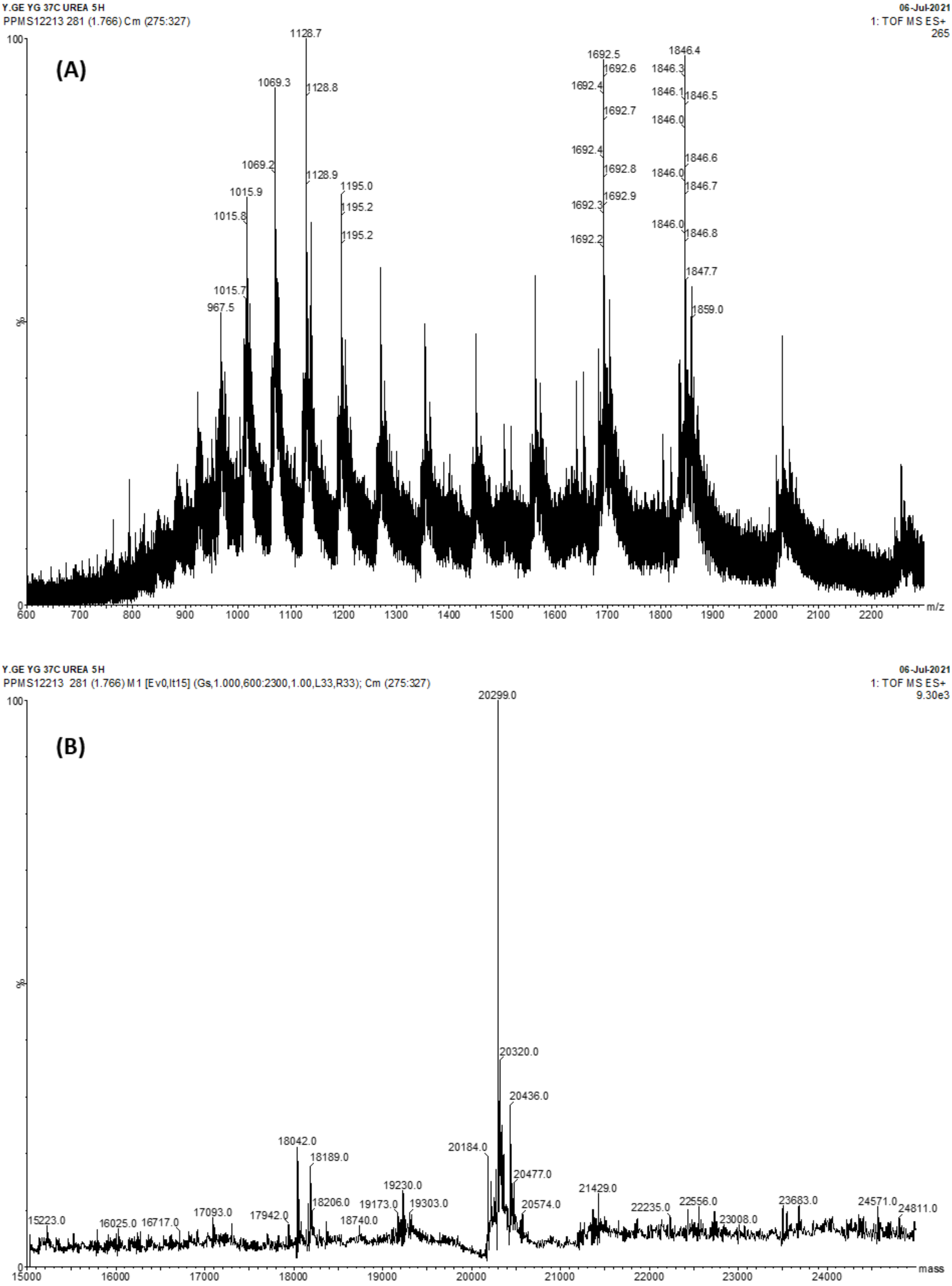


Figure S3. LC-ES mass spectra (A) and deconvolution data (B) of SD-Aβ40S26pC. The calculated mass (minus the N-terminal methionine) is 20302 and the observed deconvoluted mass is 20299.

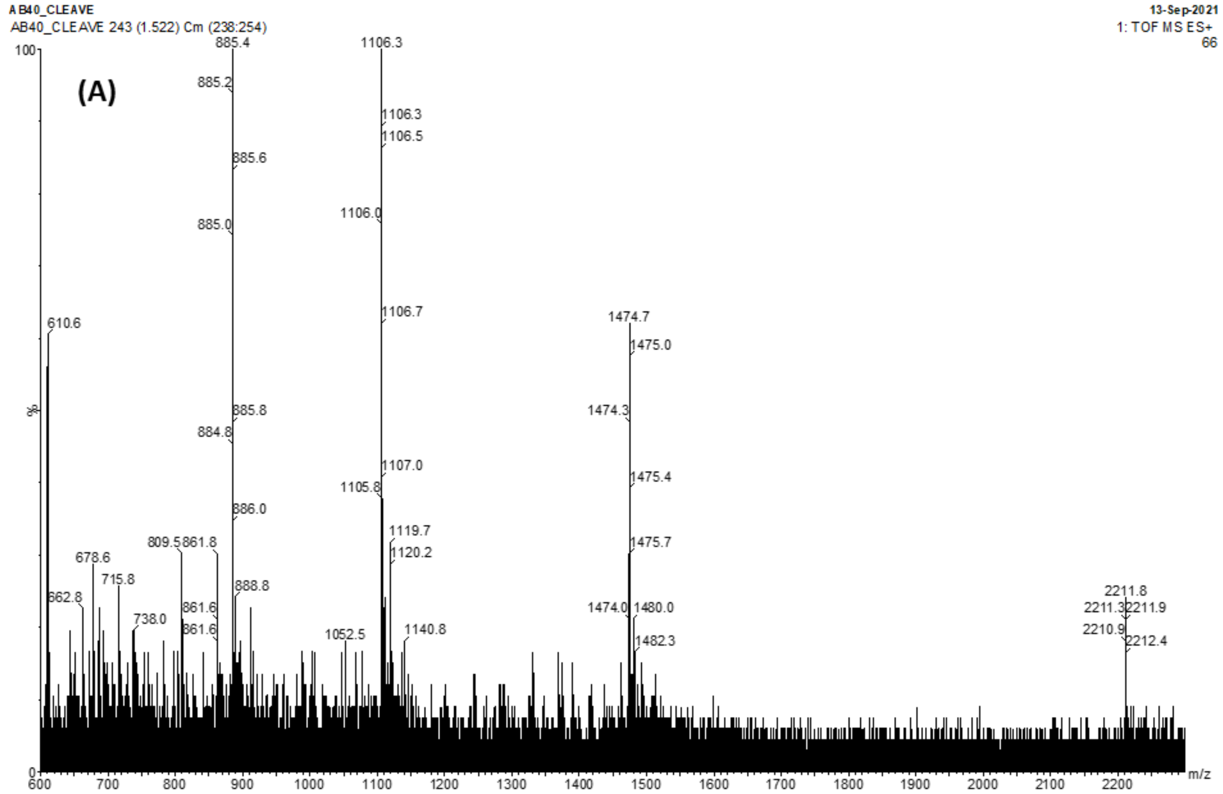


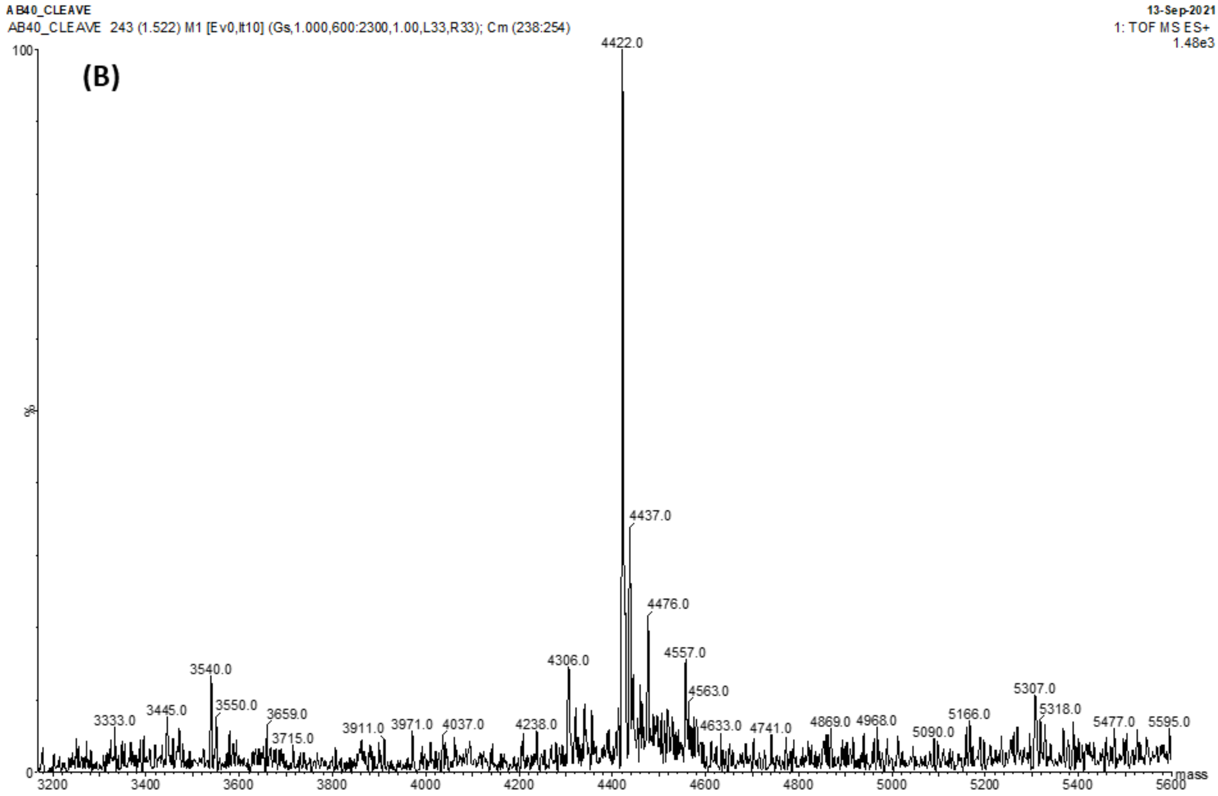


Figure S4. LC-ES mass spectra (A) and deconvolution data (B) of Aβ40-S26pC after TEV cleavage and size-exclusion chromatography. The calculated mass (minus the N-terminal methionine) is 4423 and the observed deconvoluted mass is 4422.

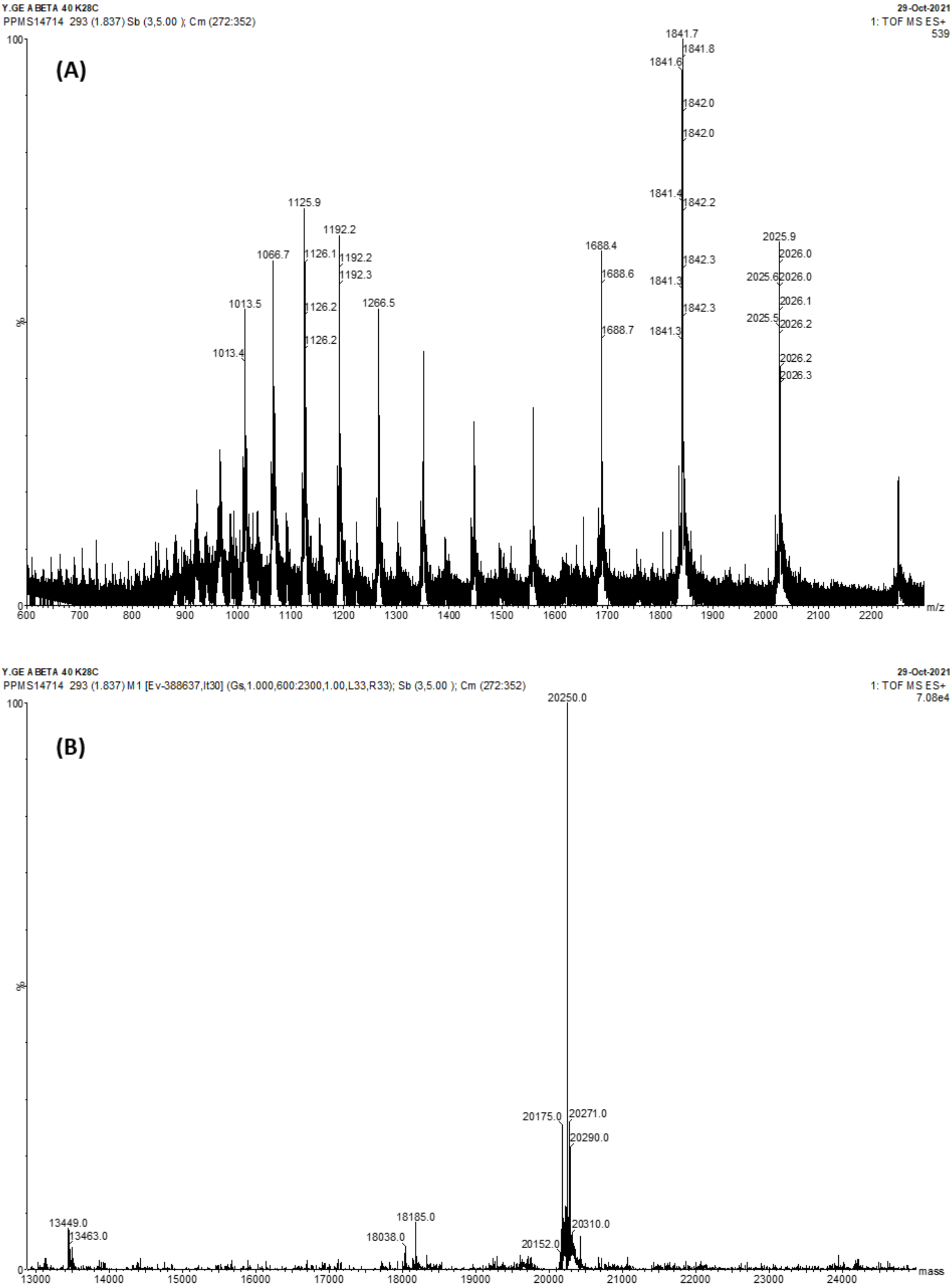


Figure S5 LC-ES mass spectra (A) and deconvolution data (B) of SD-Aβ40-K28C. The calculated mass is 20181 and the observed deconvoluted mass is 20175. There is an additional peak at 20250, which may be a BME adduct due to our purification procedure and is absent once the cysteine is converted to Dha.

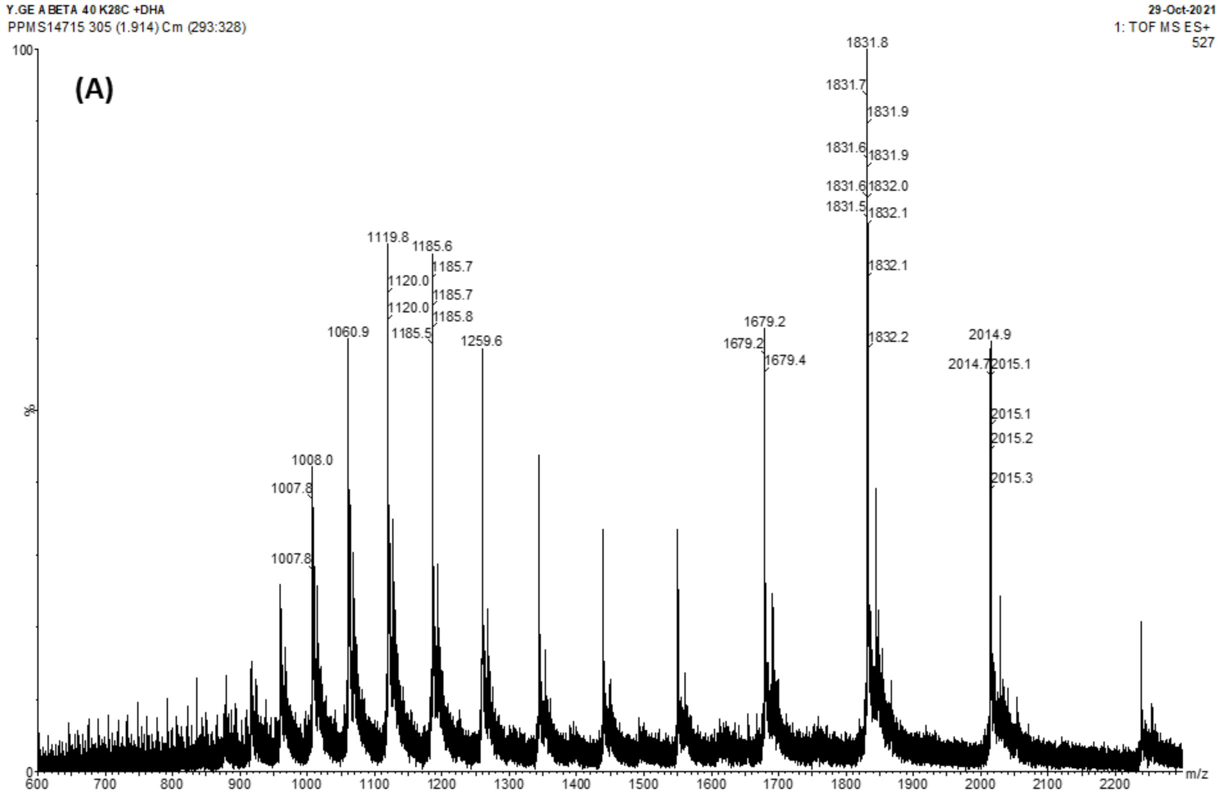


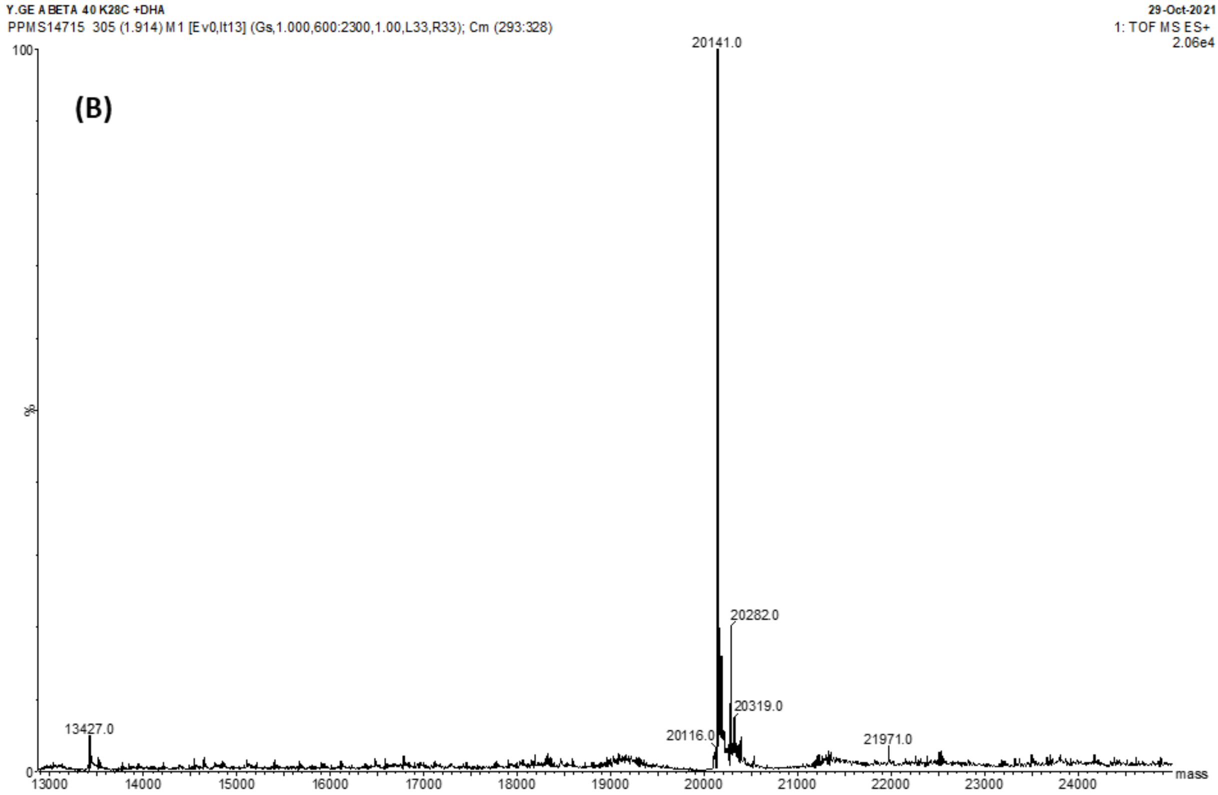


Figure S6 LC-ES mass spectra (A) and deconvolution data (B) of SD-Aβ40-K28Dha. The calculated mass (minus the N-terminal methionine) is 20147 and the observed deconvoluted mass is 20141.

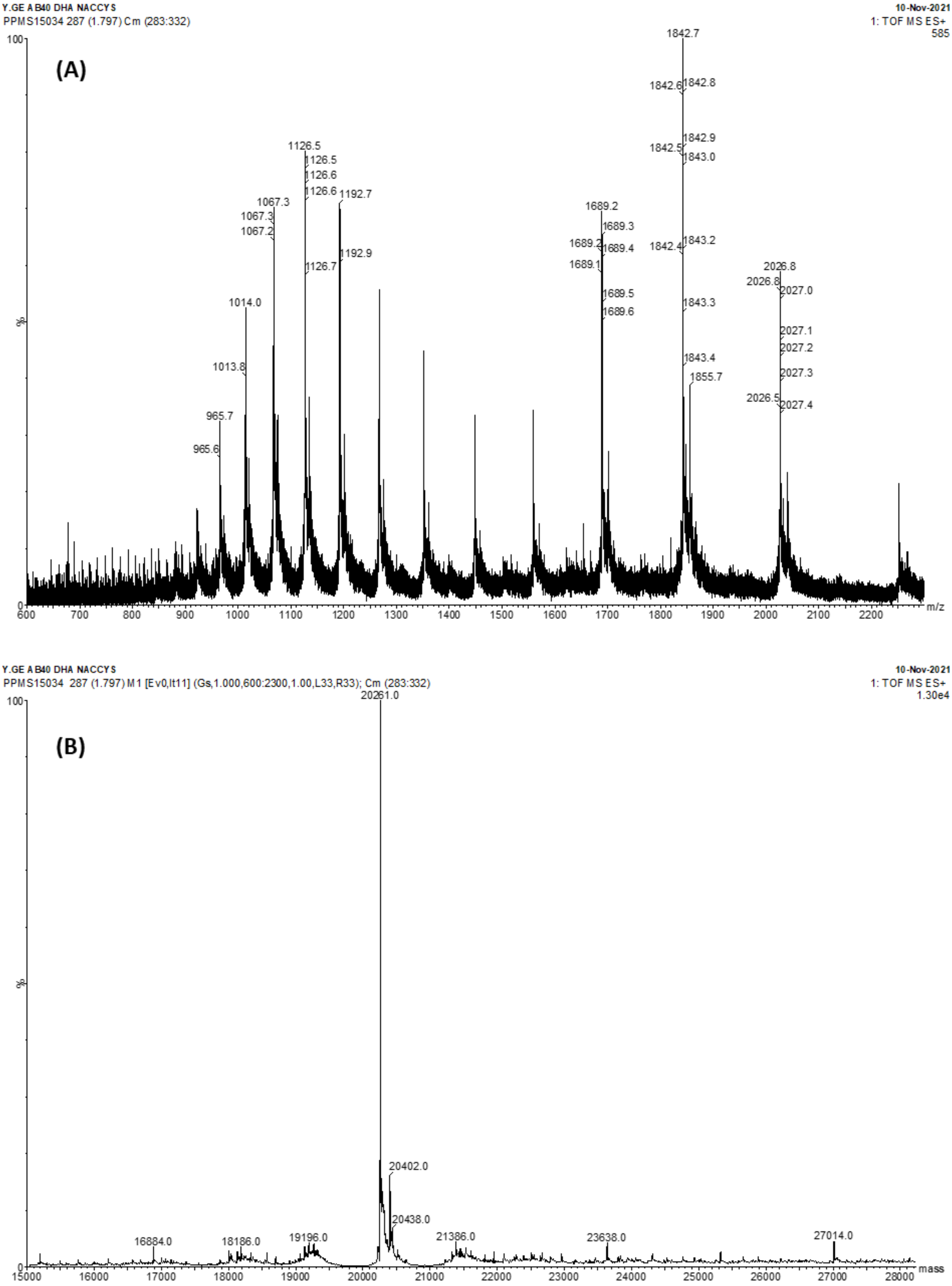


Figure S7 LC-ES mass spectra (A) and deconvolution data (B) of SD-Aβ40-K28Ac. The calculated mass (minus the N-terminal methionine) is 20265 and the observed deconvoluted mass is 20261.

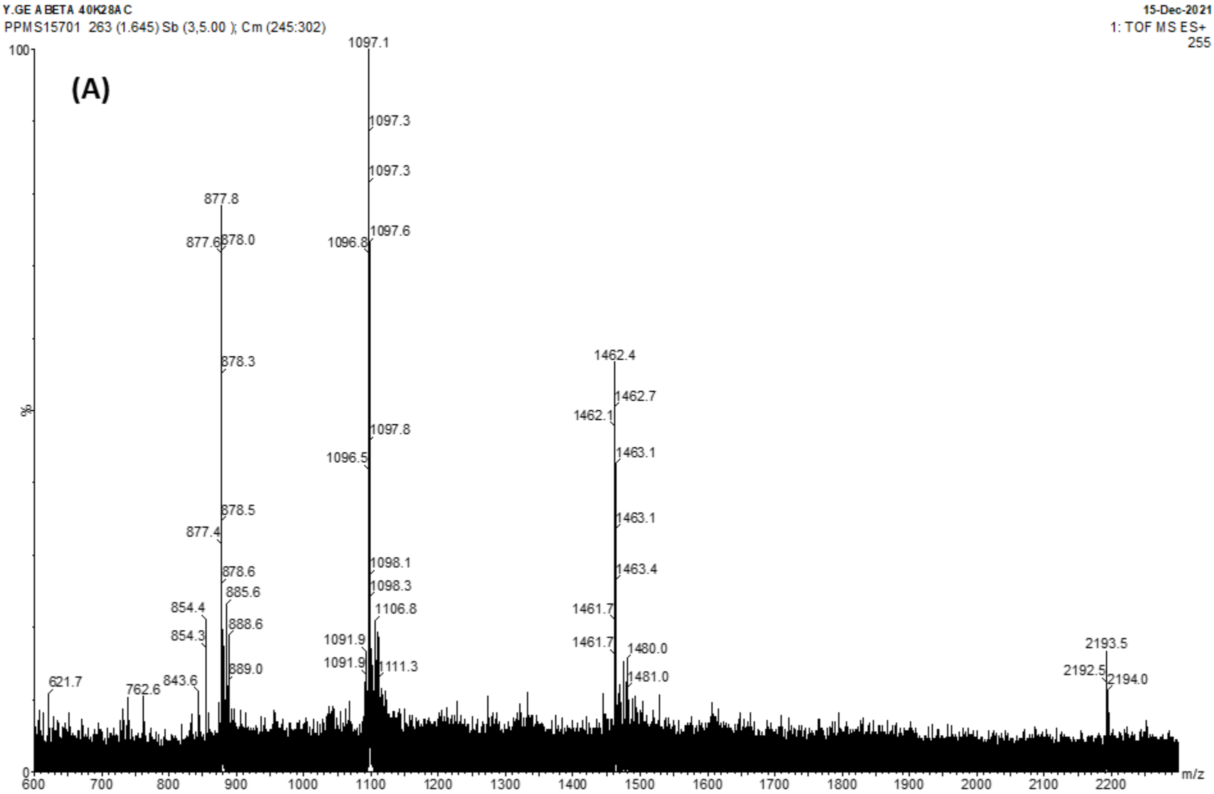


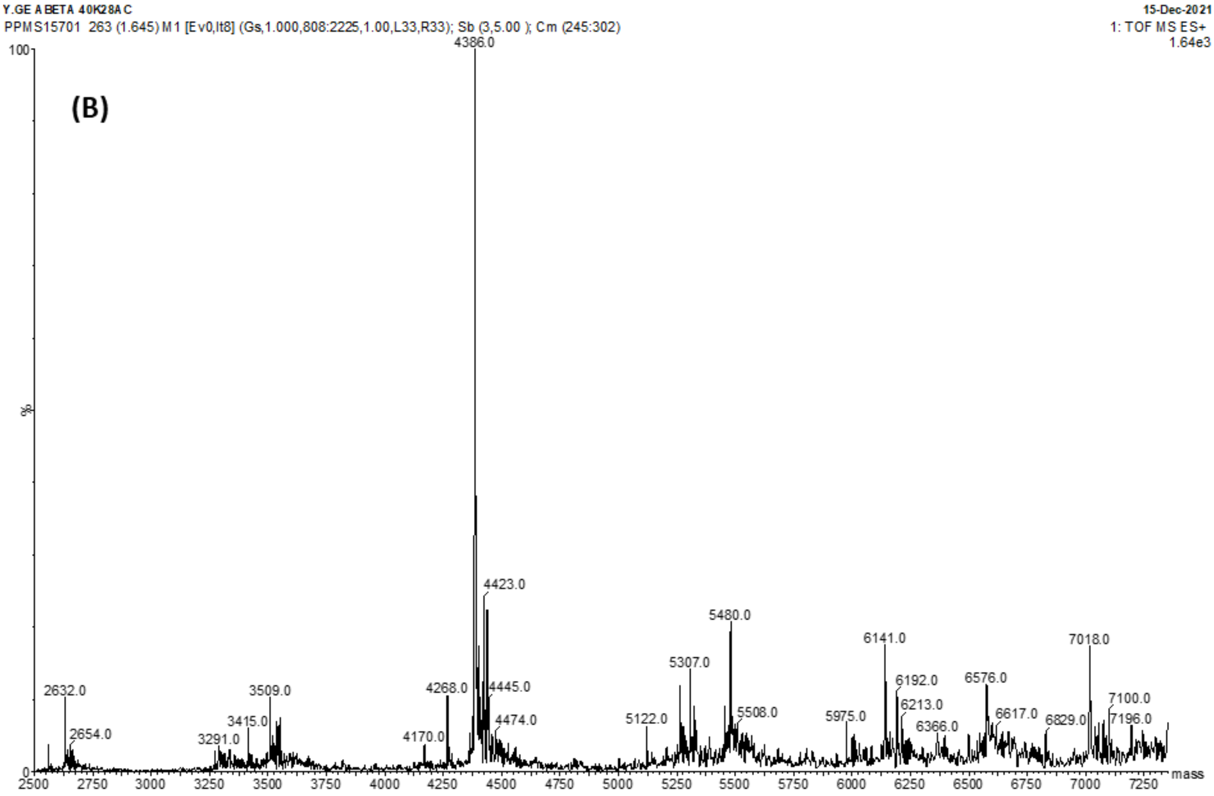


Figure S8 LC-ES mass spectra (A) and deconvolution data (B) of Aβ40-K26Ac after TEV cleavage and size-exclusion chromatography. The calculated mass (minus the N-terminal methionine) is 4386 and the observed deconvoluted mass is 4386.
